# Environmental effects on reproductive performance in preindustrial women

**DOI:** 10.1101/2023.08.06.552185

**Authors:** Lidia Colejo-Durán, Fanie Pelletier, Lisa Dillon, Alain Gagnon, Patrick Bergeron

**Affiliations:** Département de Biologie, Université de Sherbrooke, Sherbrooke, Québec, Canada; Department of Biology and Biochemistry, Bishop’s University, Sherbrooke, Québec, Canada; Department of Demography, Université de Montréal, Montréal, Québec, Canada

**Keywords:** silver spoon hypothesis, predictive adaptive response, developmental constraints, reproductive tenure, dispersal, *Homo sapiens*

## Abstract

Early life environments can have long-lasting effects on adult reproductive performance, but disentangling the influence of early and adult life environments on fitness is challenging, especially for long-lived species. Using a detailed dataset spanning over two centuries, we studied how early and adult life environments impact reproductive performance in preindustrial women. We assessed the effect of environmental conditions associated with the parish of birth and the parish where the women had their offspring. We considered resource availability differences between rural, urban, northern, and southern parishes by comparing women who switched environments during their lifetime with those who did not. We found that urban-born women had an earlier age at first reproduction and lower offspring survival to adulthood than rural-born women, while South Québec women had more offspring born than North Québec women. Moreover, switching from urban to rural led to higher offspring survival, while the reverse had the opposite effect. In addition, moving from South to North resulted in fewer offspring born and surviving, whereas moving from North to South had the inverse effect. Finally, women who changed from rural to urban and from South to North had their first child at an older age compared to those who stayed in the same urbanity type. Our study underscores the complex and interactive effects of early and adult life environments on reproductive traits, highlighting the need to consider both when studying environmental effects on reproductive outcomes.

## Introduction

Environmental effects on phenotypes are particularly pronounced when they occur early in life [1–3], especially because they can cause delayed carry-over effects on traits expressed later in life [4]. The influence of early life environment on individuals’ fitness is multifaceted, including inter-individual variation in the expression of traits that can impact survival and reproductive success [2,5,6]. For example, in red deer (*Cervus elaphus*), warm springs improve foraging conditions; cohorts born in those conditions reproduce earlier and have higher survival rates as adults compared to cold spring cohorts [7]. Similarly, semi-captive female elephants *(Elephas maximus*) born during stressful months, characterised by heavy workload, experienced faster reproductive senescence and lower reproductive success than those born during non-stressful months [8]. To measure the importance of early life environmental effects on life trajectories, it is important to quantify the direction and magnitude of these effects for a set of traits that are closely related to fitness [9,10].

There are several hypotheses proposed to explain the expected effects of early and adult life environments on fitness traits. The main ones are the silver spoon [11–13] (also known as “developmental constraints” [14]), the Predictive Adaptive Response (PAR) [15] and the quality of the adulthood environment [11]. Expectations regarding the effect of the environment on fitness for each of these hypotheses are shown in Table 1. The silver spoon hypothesis suggests that individuals born in favourable environments should perform better later in life than those who experienced poorer environmental conditions [16]. However, recent studies suggested that silver spoon effects can be non-linear, being the strongest when the adult life environmental quality is intermediate, while most individuals should perform poorly in harsh environmental conditions or do very well in excellent conditions [17,18]. Alternatively, the PAR hypothesis predicts that a match between early and adult life environment would maximise fitness, irrespectively of the environmental quality [15,19,20]. Yet, empirical support to the PAR hypothesis remains scarce in long-lived wild species [21]. Lastly, the fitness of individuals can also be influenced by the quality of their adulthood environment, regardless of their early life environment [22–24]. Therefore, it is crucial to integrate information on the quality of the adulthood environment, especially the reproductive environment, when studying the effects of early life environment on fitness [11,17,25–27]. Thus, the three hypotheses suggest a link between reproductive performance and environmental conditions, with context-specific long-term impacts of both early and adult life environments [22–24]. Notably, these hypotheses are not always mutually exclusive, but rather context-dependent, enabling us to examine which one is operative in the case of pre-industrial humans [11,18].

**Table 1:**
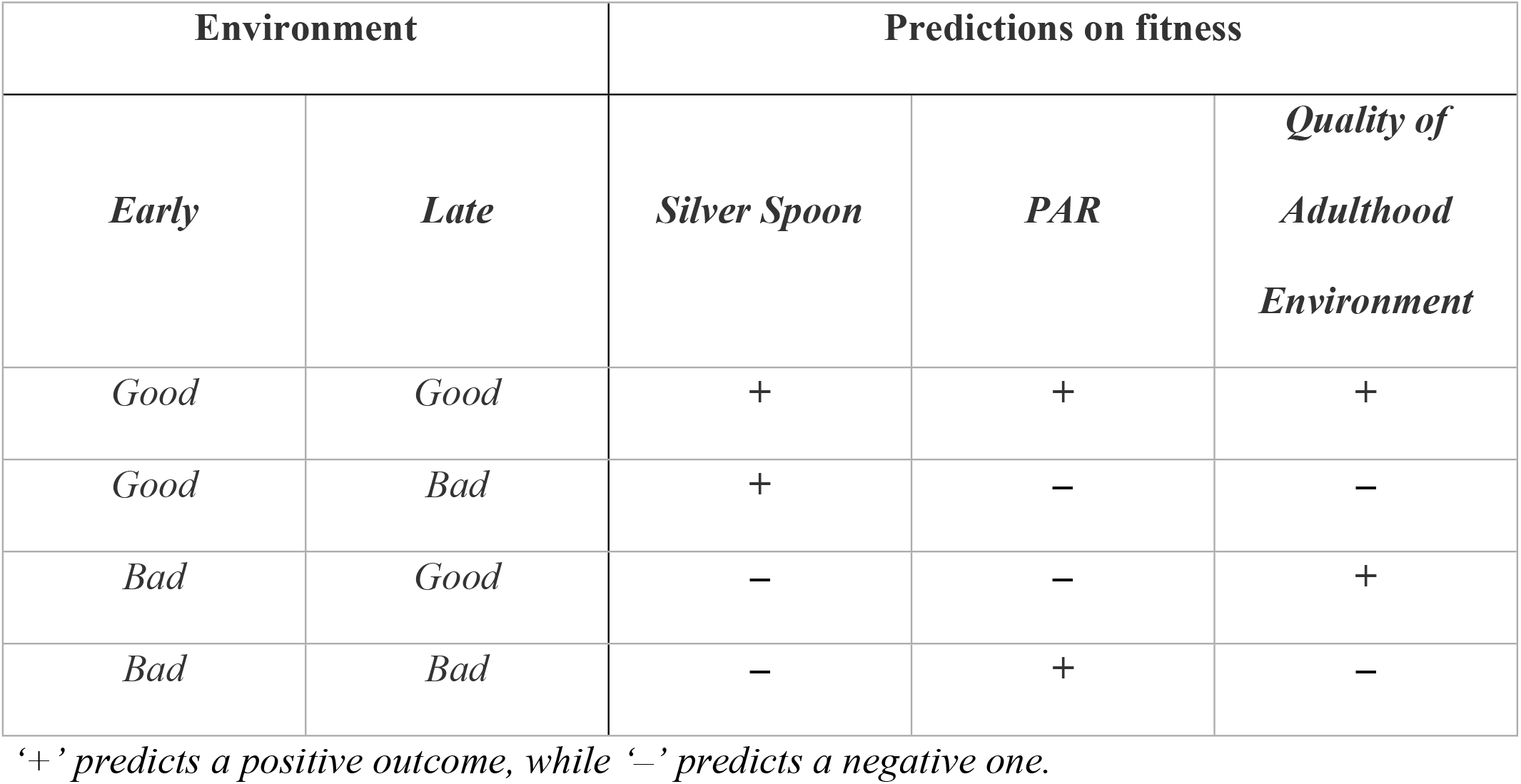
Predictions on fitness based on early and late life environmental conditions, as expected by the hypotheses of Silver Spoon, Predictive Adaptive Response (PAR), and Quality of Adulthood Environment.

| Environment |  | Predictions on fitness |  |  |
| --- | --- | --- | --- | --- |
| *Early* | *Late* | *Silver Spoon* | *PAR* | *Quality of Adulthood Environment* |
| *Good* | *Good* | + | + | + |
| *Good* | *Bad* | + | – | – |
| *Bad* | *Good* | – | – | + |
| *Bad* | *Bad* | – | + | – |

‘+’ predicts a positive outcome, while ‘–’ predicts a negative one.

To assess the influence of early and adult life environments on reproductive performance, studies typically compare individuals born in different seasons or years while living in the same location [7,8,13,28,29]. Although temporal changes in environmental conditions are of great interest [28,30], they are likely to be less pronounced compared to the habitat changes experienced by individuals that disperse. However, obtaining data on animal dispersion and its impact on lifetime reproductive success in wild populations is challenging due to the need for comprehensive longitudinal data on reproduction throughout the entire lifespan [31].

Long-term datasets on preindustrial human populations are very valuable to disentangle the effect of early and adult environments on fitness, as they generally provide complete life histories of several hundred individuals over very long periods (>100 years [32]). Parish registers have detailed individual-level information on family size, genealogy, and lifespan, and have been widely used to study life history patterns of preindustrial populations to help answer evolutionary ecological questions [33]. Parish registers also provide the location where major life history events happened, allowing to track movement of individuals over long distances throughout their life [34], which can rarely be done with wild animals over the long term. One of the largest databases on preindustrial humans comes from the *Registre de la population du Québec Ancien* (RPQA) [35]. This dataset covers about the first two centuries of settlement in Nouvelle-France (current Province of Québec, Canada) and includes almost half a million individuals with reconstructed life history over nine generations [35], making it an excellent resource for studying environmental effects on fitness.

Here, we investigated the relative importance of early and adult life environments in shaping lifetime reproductive performance of preindustrial women. Previous studies have shown that rural parishes and the south shore of the Saint Lawrence River offered better environmental conditions than the urban parishes and the north shore, such as greater food availability [29,36,37] (See Fig. S1). To separate the relative effects of early and adult life conditions, we only considered women who changed parish between their birth and the birth of their offspring. In this dataset, some women relocated but remained in a similar environment while others relocated and switched urbanity and/or shore, allowing us to disentangle the effect given by their early life environment from the effect given by their environmental switch in adulthood. We also aim at investigating whether the environmental effects on reproductive performance are better explained by the silver-spoon hypothesis, the PAR, or the environmental quality during adulthood. We observed that all the analysed traits exhibit potential involvement of the three hypotheses.

## Materials and methods

### Study population and historical context

The overall RPQA database includes 448,501 individuals and 74,067 families [35], from 1572 (earliest date of birth of a settler, before migrating to the colony) to 1870 (latest date of death of an individual) in 166 different parishes (Fig. S1). It was assembled by reconstituting family histories from Roman Catholic parish registers of baptisms, marriages, and burials from which the date and location of the events are known [35,38,39]. We filtered this database to only keep women who were born in Nouvelle-France, with known parents, with known date and location of birth and death (n = 216,442), and who did not remarry (93.6% of women). As we were interested in contrasting early and adult life environments, we also only included women who had relocated to a different parish between their birth and reproduction. In addition, we selected two subsets, one with women born between 1640 and 1750, which contains 7,203 women who had 61,606 offspring, of whom 29,059 survived to adulthood, and the other with women born between 1640 and 1729, which contains 3,959 women who had 33,878 offspring, of whom 19,865 survived to adulthood (see Fig. S2 for more details on filtering). This filtering ensured that all the women expressed a complete reproductive history, since the date of death of the descendants is less precise at the end of the record (see also [34]). For further descriptive statistics of the subsets, see Tables S1, S2 and S3.

During the study period, there was limited emigration from the colony, such that the population is considered semi-closed [35,38–40]. The population was composed primarily of farmers, with a smaller number of artisans, merchants, civil servants, professionals, and elites residing in urban areas [41]. Québec City (founded in 1608) and Ville-Marie (founded in 1642, now known as Montréal) were the only urban parishes along the St. Lawrence River [41–43]. Nouvelle-France experienced lower overall mortality rates compared to Europe at the same time [44–46], although infant mortality rates were similarly high [47]. The population was characterised by a high fertility rate compared to other preindustrial human populations [35], which led to exponential growth [48,49], with the average family size in our subset being 8.6 (Table S1). Extramarital relationships were socially condemned and very rare; extra-conjugal paternity has been estimated to be less than 1% [50,51].

### Environmental conditions

The environmental characteristics of the area were distinctive, with the north and south shores of the St. Lawrence River differing in average temperatures, persistence of snow cover, and soil fertility [52]. For example, the frost-free season was about one month longer in the south-west than in the north-east of the study area [29]. During the 1688 census, it was reported that the south had a yield of 14.5 bushels (511 litters) of grain per inhabitant from the previous fall’s harvest, while the north had a yield of 10.7 bushels (377 litters) per inhabitant over that year [29]. Consequently, inhabitants in the southern part of the St. Lawrence River Valley experienced better agricultural conditions and their farms were more productive than those from the North. In addition, there was a differential mortality between urban and rural populations due to socio-economic factors [41], given that rural inhabitants had greater food availability, lower exposition to diseases and lower infant mortality than urban populations [36,48]. Epidemic cycles were also common in both Québec City and Montréal because of the increasing urban population density and poor hygiene [46,53]. Women who lived in Québec City and Montréal were thus classified as urban residents and the rest of the population was designated as rural, with urban parishes being considered a worse environment than rural parishes.

The early life environment was characterised by the variable *Birth Environment,* which had four levels: Rural South (both good environments), Rural North (good shore, bad urbanity), Urban South (bad urbanity, good shore) and Urban North (both bad environments). To assess the impact of switching environments between early life and adulthood, we used two variables: *Switching Urbanity* and *Switching Shore*. The variable *Switching Urbanity* indicated whether women stayed in the same urbanity while *Switching Shore* indicated whether they stayed on the same shore. Thus, both variables had three levels, representing the same environments or transitions from bad to good or good to bad environments. We also considered the effect of the distance between the early and adult life parishes, as it has already been shown to influence fitness [34]. We calculated it as the Haversian distance (km) between the geographic coordinates of the parish of birth and the parish of first reproduction, using the *geosphere* package [54].

### Reproductive performance

We used three main traits to quantify women’s reproductive performance: age at first reproduction (AFR), defined as the age at which a woman first reproduced, number of offspring (NO), corresponding to the number of children born to each mother, and lifetime reproductive success (LRS), corresponding to the number of infants who survived to at least 15 years of age, considered to be the age of maturity in preindustrial human populations (e.g. [34,55]). Survival to 15 was established by the presence or absence of subsequent life events when the date of death was unknown [34]. A better reproductive performance would be given by a younger AFR [32,56] and a higher NO and LRS.

We also considered two additional traits: the *fertile years*, which is the number of years during which a woman had the opportunity to reproduce, and the ratio between LRS and NO (LRS/NO), which estimates the proportion of offspring who survived to adulthood. We accounted for the number of *fertile years* (also called reproductive tenure [57,58]) on AFR, NO and LRS, but also included it as response variable in a model that we report in the supplementary material (Table S4 & S5, Fig. S3A & S4A). Additionally, since the results for LRS/NO were qualitatively similar to the result for LRS (see below), they are reported in the supplementary material (Table S4, Fig. S3B & S4B, C) and we do not discuss them further. We also performed a sensitivity analysis on the age at marriage, which is the difference between the date of birth and the date of the wedding (Table S4, Fig. S5), to ensure the accuracy of our measurements for AFR, and reported it in the supplementary material.

### Statistical analyses

We ran three sets of models, one for each reproductive trait. AFR was normally distributed and analysed with a linear mixed model (LMM). NO and LRS were analysed with a generalised linear mixed model (GLMM) with a Poisson distribution. The models accounted for early life environment with the variable *Birth Environment* and for environmental switch between birth and adulthood with the variables *Switching Urbanity, Switching Shore,* and the interaction between them. Other fixed effects included in the model were *fertile years* and *wave front*. *Fertile years* was only included in the models for NO and LRS and was calculated as the number of years between the date of marriage and death of either spouse, or 45 years old when both partners were still alive [34]. *Wave front* has been shown to influence family size [59] and was calculated as the difference between a life-history event (first reproduction in our case) and the date of the foundation of the parish of birth. All models included the random terms *family identity* (marriage ID of the parents of the woman), to control for genetic and shared family environment effects, and *year of birth*, to account for potential cohort effects [56].

All analyses were performed using R version 4.3.1 with the package *glmmTMB* [60]. We conducted likelihood ratio tests (LRT) to assess the statistical significance of the fixed terms and a stepwise backward procedure to remove the non-significant terms from the model until the final models were reached [61]. We calculated the conditional and marginal R² for the final models with the package *MuMIn* [62] and for each variable in the final model we calculated the partitioned R² with the package *glmm.hp* [63]. We also performed post-hoc Tukey tests with the package *emmeans* [64] to conduct pairwise comparisons of environmental effects on reproductive performance.

## Results

### Age at first reproduction

The best model explaining AFR included both early life environment (LRT *Birth Environment*: χ² (3) = 102.95, p <0.001) and environmental switch between birth and adulthood (LRT *Switching Urbanity* * *Switching Shore*: χ² (4) = 6.76, p= 0.149), as well as the *wave front* and *distance* (Table 2). Compared to women born in rural areas, those born in urban areas had their first child on average 3.3 and 4.6 years earlier, depending on whether they were born in the south or the north shore, respectively (Fig. 1A), and this difference was significant according to post-hoc Tukey’s pairwise multiple comparison test (hereafter called Tukey test). In addition, women born in the southern urban parishes began reproducing 1.3 years later than those born in the northern urban ones, which was also significant according to the Tukey test (Fig. 1A). For the interaction between *Switching Urbanity* and *Switching Shore*, women who remained in the same urbanity reproduced on average 1.5 years earlier, than those who switched from rural to urban areas and from south to north, which was the only combination that the Tukey test indicated as significantly different (Fig. 2). *Distance* between early and adult life environments also influenced AFR (Table 2); women who moved greater distances began to reproduce later (Fig. S6A). Overall, 14% of the variance in AFR was explained by the fixed effects of the final model (Table S5), and most of the variance explained was due to the *wave front*, followed by the early life environment (Table S5).

**Figure 1:**
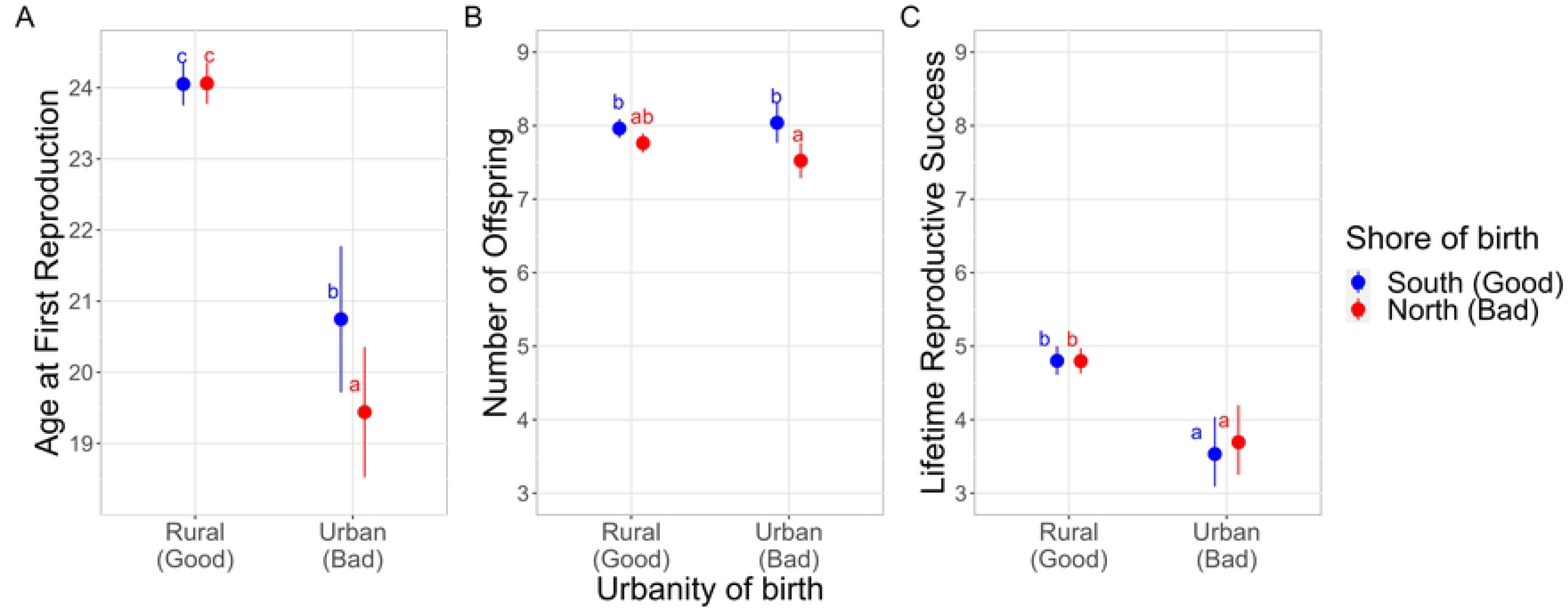
Early life environmental effects on Age at First Reproduction (panel A), Number of Offspring (panel B) and Lifetime Reproductive Success (panel C) according to the environment of birth, given by the urbanity (Rural or Urban) and the shore (North and South). Rural and South are considered good environments and Urban and North are considered bad environments. The dots are the predicted marginal values, and the lines are their confidence intervals. Estimates with different letters are statistically different, given by a post-hoc Tukey’s pairwise multiple comparison test (P <0.05).

**Figure 2:**
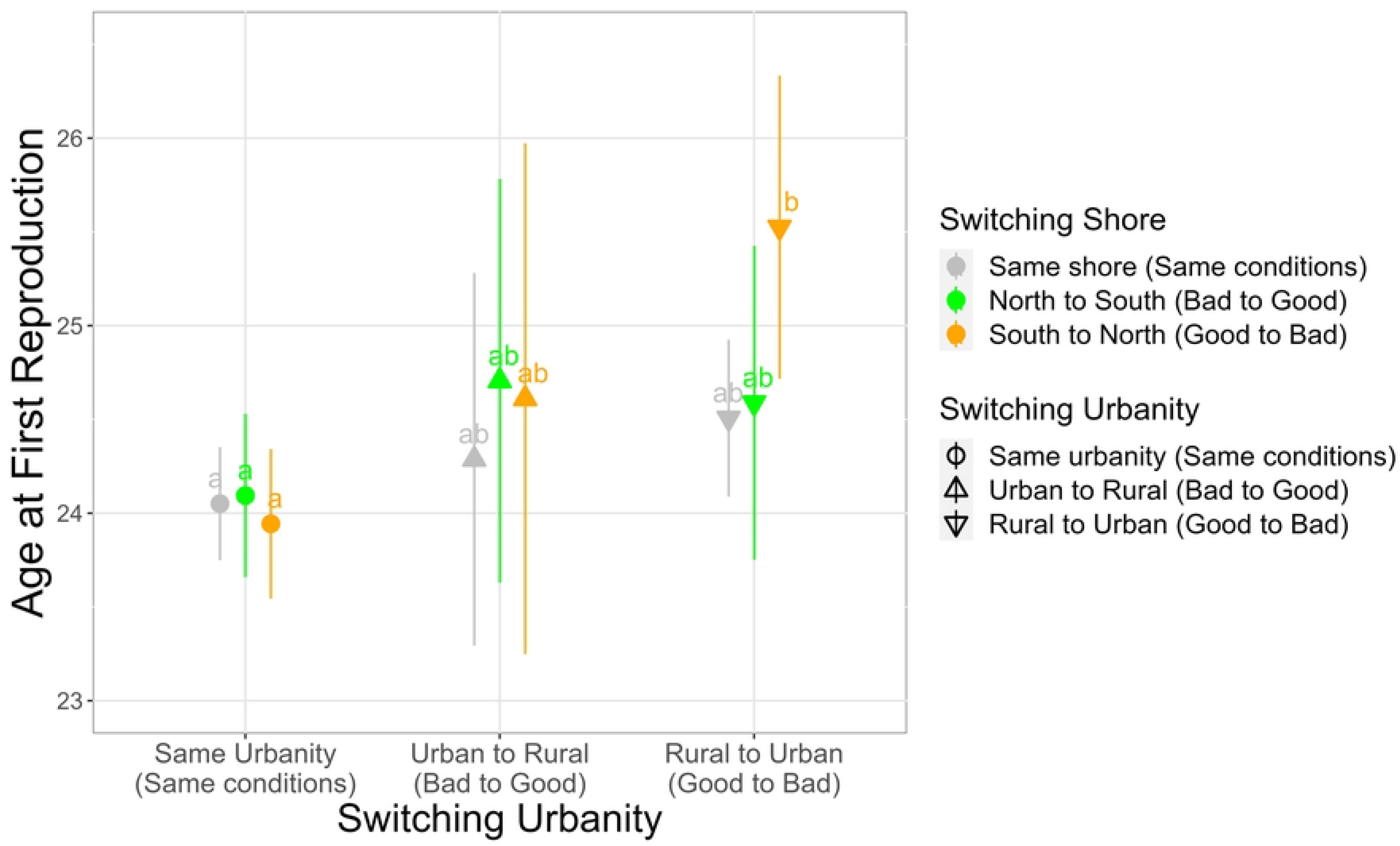
Adult life environmental effects on Age at First Reproduction according to the interaction between the switch in urbanity and the switch in shore. Staying in the same urbanity or in the same shore is considered as staying under the same conditions, while switching from urban to rural or north to south is seen as going from a bad to a good environment, and switching from rural to urban or south to north is considered going from a good to a bad environment. The dots are the predicted marginal values, and the lines are their confidence intervals. Estimates with different letters are statistically different, given by a post-hoc Tukey’s pairwise multiple comparison test (P <0.05).

**Table 2:**
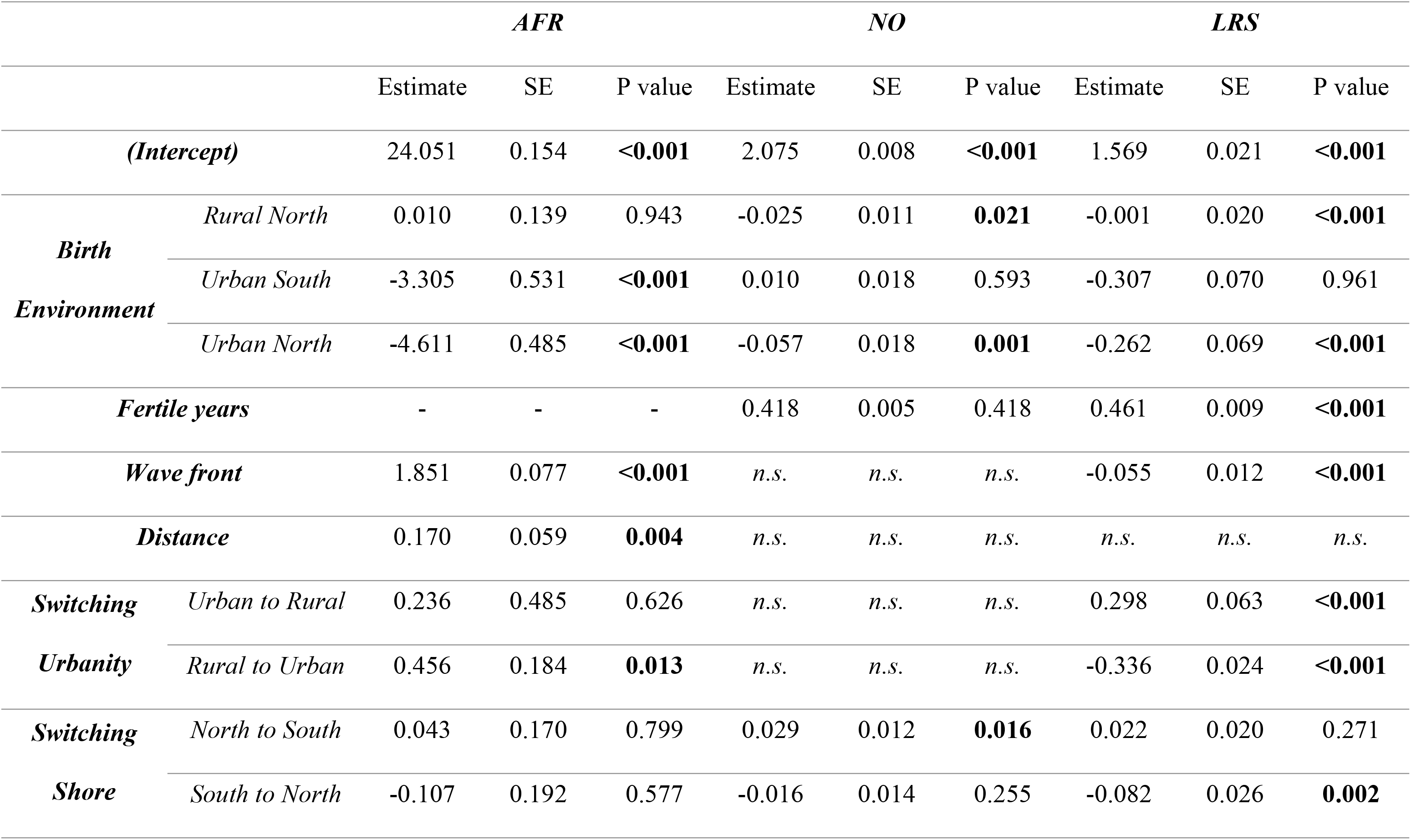
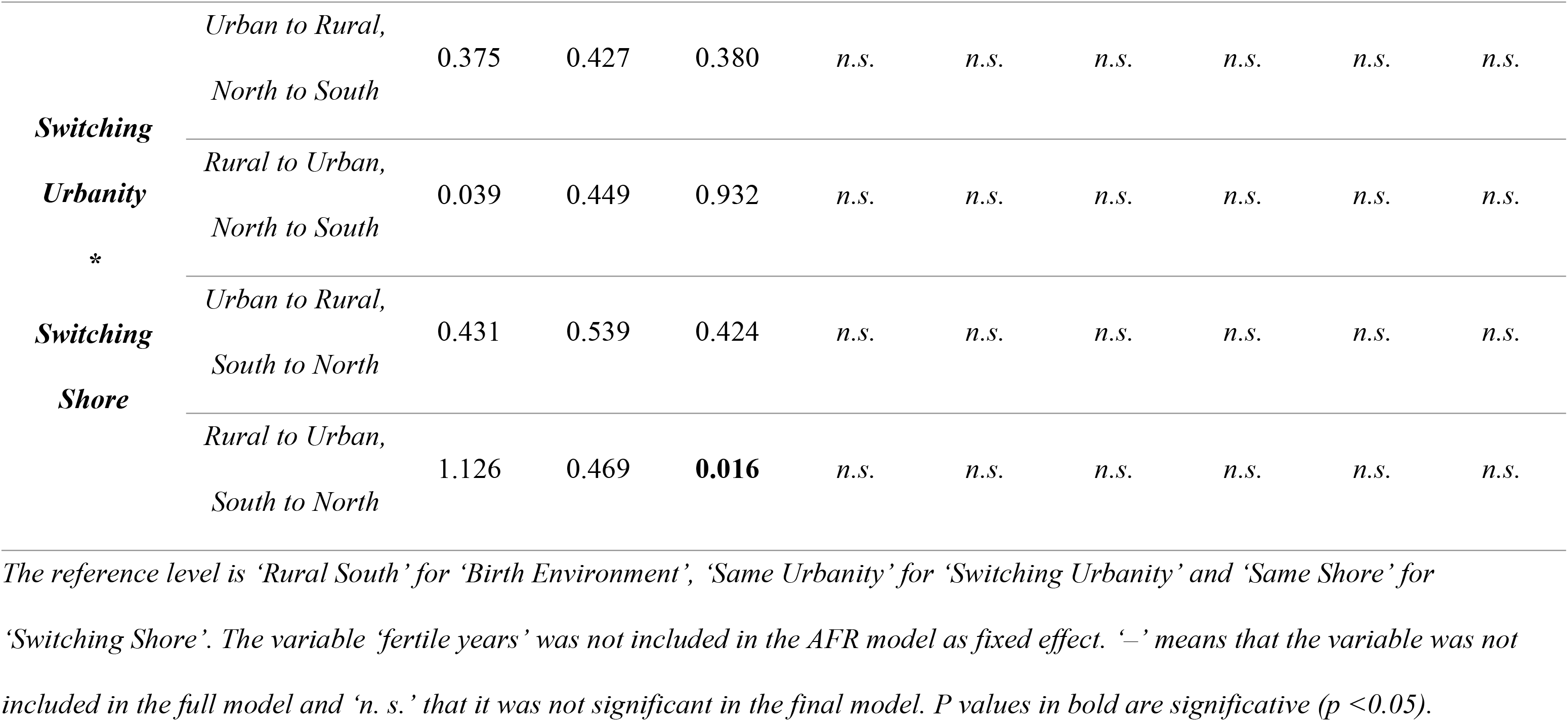
Output of the final models selected by stepwise backwards selection from a full model with all the variables present in the first column, for Age at First Reproduction (AFR), Number of offspring (NO), and Lifetime reproductive success (LRS).

### Number of Offspring

The best model explaining the number of offspring born included early life (LRT *Birth Environment*: χ² (3) = 12.69, p=0.005) and environmental switch between birth and adulthood (LRT of *Switching Shore*: χ² (2) = 7.06, p = 0.029) as well as *fertile years* (Table 2). Women who were born in parishes located south of the St. Lawrence River had a higher number of offspring compared to women born in the north (an increase of 0.2 for rural populations and 0.5 for urban populations), but according to the Tukey test this difference was only significant for the urban parishes (Fig 1B). There was also a significant difference in NO between rural south and the urban north parish, but not between rural north and both urban parishes, based on Tukey test (Fig 1B). When analysing the environmental switch before adulthood, the variable *Switching Urbanity* was not significant (LRT: χ² (2) = 3.85, p = 0.146) but *Switching Shore* was significantly related to the number of offspring born (Table 2, Fig. 3A). Women who switched from north to south (bad to good) had 0.2 more offspring than women who stayed on the same shore and 0.4 more than the ones who moved from south to north (good to bad), and the difference was significant according to the Tukey test (Fig. 3A). Although the fixed effect of this model explained a moderate amount of variation (marginal R^2^ = 0.57), most of the variation was explained by the number of *fertile years* (Table S5).

**Figure 3:**
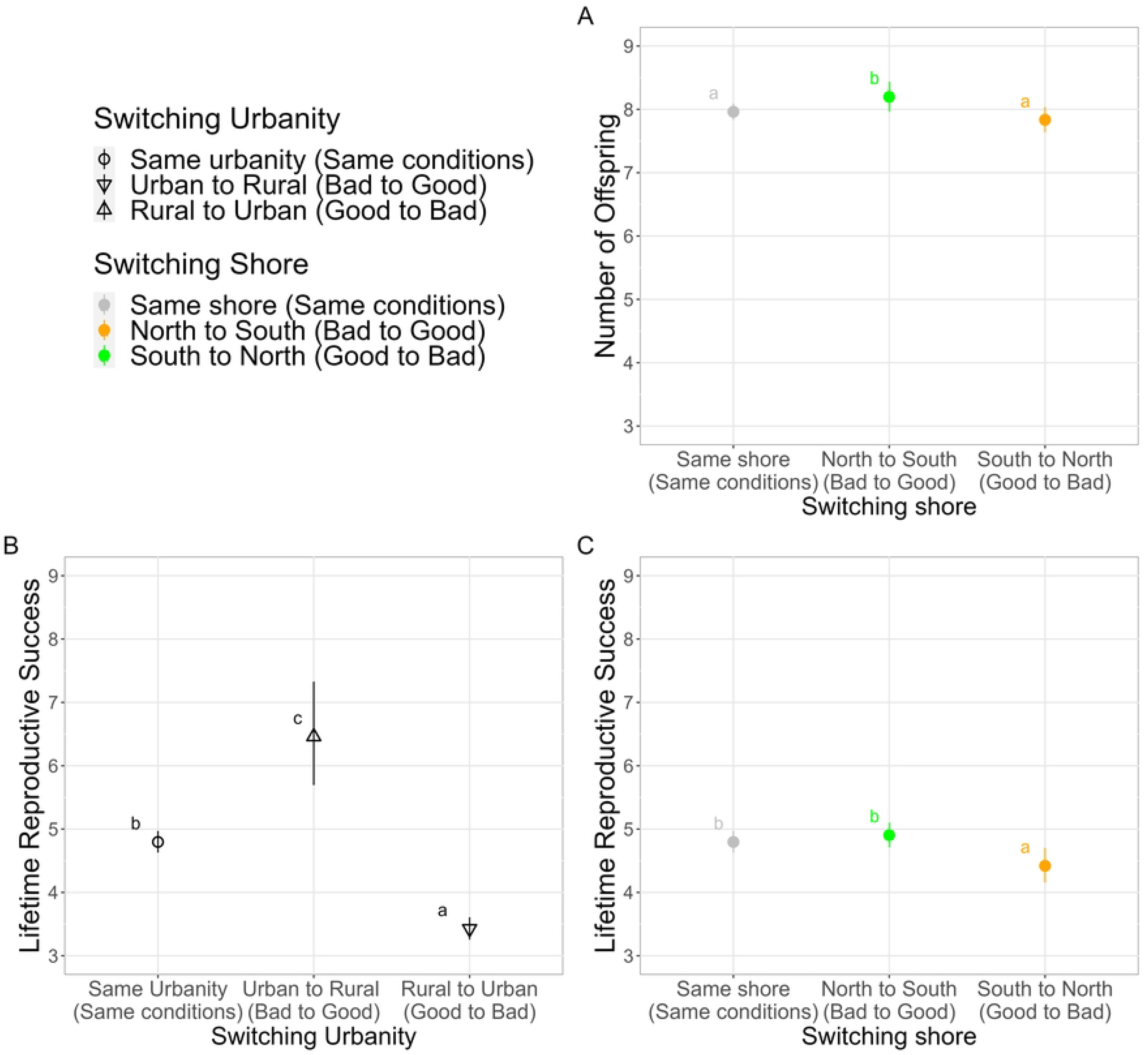
Adult life environmental effects on Number of Offspring (panel A) and Lifetime Reproductive Success (panel B & C), according to the switch in urbanity and the switch in shore. Staying in the same urbanity or in the same shore is considered as staying under the same conditions, while switching from urban to rural or north to south is seen as going from a bad to a good environment, and switching from rural to urban or south to north is considered going from a good to a bad environment. The dots are the predicted marginal values, and the lines are their confidence intervals. Estimates with different letters are statistically different, given by a post-hoc Tukey’s pairwise multiple comparison test (P <0.05).

### Lifetime Reproductive Success

The best model explaining the lifetime reproductive success included early life environment (LRT *Birth Environment*: χ² (3) = 21.35, p <0.001) and environmental switch between birth and adulthood (LRT *Switching Urbanity*: χ² (2) = 238.04, p <0.001; LRT *Switching Shore:* χ² (2) = 11.31, p =0.003) as well as *fertile years* and *wave front* (Table 2). Women born in urban parishes had on average 1.2 less offspring surviving to adulthood than the ones born in rural parishes, which was significant according to the Tukey test (Fig. 1C). However, there was no significant difference in LRS between women born in the south shore compared to the north shore (Fig. 1C). Women who switched from rural to urban (good to bad) had 1.4 less offspring surviving to adulthood than women who stayed under the same conditions and 3 less than women who switched from urban to rural (bad to good), and the difference was significant according to the Tukey test (Fig. 3B). Therefore, women who switched from urban to rural (bad to good) had 1.7 more offspring surviving than the ones that remained in same conditions (Fig. 3B). Women who switched from south to north (good to bad) had on average 0.4 less offspring surviving to adulthood compared to the ones who stayed in the same shore or who moved from north to south (bad to good), which the Tukey test indicated as significant (Fig. 3C). The marginal R^2^ of the final model was 0.53 and most of the variation was explained by the number of *fertile years*, followed by whether a woman switched urbanity and the *wave front* (Table S5).

## Discussion

By capitalising on data from women who relocated parishes during their life, we were able to disentangle the effects of early life and adulthood environments on reproductive performance. Given that in our dataset all women relocated parishes between their birth and their reproductive events, we controlled, at least partially, for the cost of migration. Our study shows that both the early and adult life environments affected reproductive performance, but the magnitude of these effects differed depending on the trait. Our results offer support for the silver spoon hypothesis for two fitness proxies (NO and LRS), while the effect on AFR and LRS support the expectations of the PAR hypothesis. The adulthood environment had effect on all the studied traits.

The silver spoon hypothesis states that individuals born in a good environment will systematically have a better performance than those born in a bad environment ([11–13], Table 1). We found that women born in southern parishes had a larger NO and that women born in rural parishes had a larger LRS. These results are consistent with further findings from other preindustrial populations like the Finnish, for whom better early-life nutrition was associated with improved reproductive performance [65]. Furthermore, it has been observed that increased malnutrition and higher prevalence of infections during women’s early life results in a heightened rate of neonatal mortality among offspring of stunting mothers [66], which has also been observed in some populations of wild animals [67]. Different nutrition and environmental factors during early life affect distinctly to the oocyte and embryo development, prenatal survival, and offspring number and quality [68], which may explain why we observe different environmental characteristics (such as shore or urbanity) showing distinct silver spoon effects in the traits used to characterise reproductive performance (such as NO or LRS).

In our analyses, we also found evidence in favour of the quality of the adulthood environment in all the examined reproductive traits, and evidence for PAR in AFR and LRS. As previously mentioned, according to the PAR hypothesis, it is anticipated that individuals who have a matching environment in adulthood compared to their early life environment will exhibit better performance [15,19,20], whereas the hypothesis regarding the quality of the adult life environment posits that individuals living in a favourable environment during adulthood should display better performance irrespective of their early life circumstances [11,17,25–27]. Due to limitations in our available data, we are unable to fully separate the specific effects attributed to PAR from the ones explained by the quality of the adulthood environment (Table 1). For AFR, staying in the same urban environment led women to start reproducing earlier than those who switched urbanity and shore, supporting predictions from PAR [15,19,20]. Mechanisms allowing anticipatory matching may have aided ancestral survival amid environmental changes, favouring early reproductive success but compromising survival later in life [15]. Besides, among the evidence that supported the hypothesis of the quality of the adulthood environment [17], we observed that only women who moved to the worst environment experienced a detrimental effect on AFR. Age at marriage showed a consistent pattern of results in line with the findings for AFR. Although our results for AFR support both the PAR hypothesis and the importance of the quality of the adulthood environment (Table 1), the effect size of the environmental switch was relatively smaller when compared to the effect observed for the early life environment, highlighting the significance of the early life environment as well as the relevance of the silver spoon hypothesis in this context.

The transition from the south to the north shore is the only one that negatively impacts both NO and LRS, which is consistent with previous findings regarding the detrimental effects of switching shore on other traits (such as longevity) on our population [37]. For LRS, since the match between the early and the adulthood environment and the switch from a bad to good environment brought the same positive outcome, it is not possible to disentangle the comparative relevance of PAR and the importance of the quality of adulthood environment (Table 1). Nonetheless, staying on the same shore is equally detrimental for NO, which favors the hypothesis of importance of the quality of the adulthood environment. Regarding the switch in urbanity, higher LRS is achieved by women who switched from urban to rural environments. Given the better sanitary conditions found in rural parishes [46,53], this outcome highlights how the quality of the adulthood environment is crucial for offspring survival, reflecting epidemiological evidence in human populations linking poor maternal nutrition during conception or early pregnancy to increased rates of premature birth and offspring health issues [25,69,70]. The importance of the adulthood environment from the perspective of the mother is difficult to distinguish from the importance of the early life environment from the perspective of the offspring, an idea advocated by the silver spoon hypothesis that is also consistent with the greater child mortality previously observed in urban settings in our population [47]. Furthermore, staying in the same urbanity led to a higher LRS than switching from rural to urban, which provides additional support to the PAR hypothesis. Previous studies on the Dutch famine found no evidence of PAR on fertility, but they compared different conditions given by year of birth and not by the early life environment, as in our case [71]. Since natal dispersal effects on reproductive success can vary with phenotypes and environments, it is not surprising that we find diverse results for urbanity and shore [72]. Therefore, our findings regarding LRS support both the PAR hypothesis and the quality of the adult life environment and our findings for NO favor the latter. However, the early life environment demonstrates a similar effect size to the switch in shore for NO, surpasses the effect of the switch in shore for LRS, and is smaller than the effect size of the switch in urbanity for LRS. Consequently, the quality of the adult life environment provides a better explanation for the observed effects on LRS, followed by the silver spoon and PAR, due to the effect size. Additionally, a combination of silver spoon, and the importance of the quality of the adult life environment is important for NO.

Some of our results are open to multiple interpretations, as they can be explained by distinct hypotheses. For example, we found that women born in urban parishes give birth to their first child earlier than those born in rural ones. The same pattern was observed when analyzing the age at marriage. Given that a positive correlation between earlier AFR and higher NO and LRS exists (Fig. S7), it is surprising that women born in urban parishes did not have more children nor more children surviving to adulthood than women born in rural parishes. A correlation between a younger AFR and higher LRS was also reported on a small population at the Ile aux Coudres (QC, Canada) [56,73] and in a preindustrial Finnish population a negative correlation between earlier AFR and lifetime fitness was found [74]. The lack of precise socio-economic factors for each individual due to the particularities of Catholic church registers in Québec, such as their wealth or profession, prevents us from being able to make a more comprehensive analysis of this result [35]. Nonetheless, our study could possibly align with the earlier onset of reproduction of wealthier women seen on the Finnish population [75,76], since there was a greater presence of higher income occupations among the inhabitants of Québec City and Montréal [41,77,78], which would support the silver spoon hypothesis but not our prediction that rural environments are better than urban ones. In that case scenario, life history trade-offs suggest that females who experienced early adversity may direct more energy towards growth or maintenance in their early adulthood to secure future reproductive potential [79], which could be the strategy adopted by rural-born women. Alternatively, poorer early life conditions have been related to a first pregnancy at a younger age on British women, with detrimental consequences for their offspring [80], which is in line with the poorer health and diet conditions expected in preindustrial urban environments compared to the rural ones [46]. If this was the case, the observed connection between younger AFR of urban born women and their lower NO and LRS could be a trade-off between an earlier onset of reproduction and a decline in fitness during later life stages [81,82].

To improve our understanding about the indirect impacts of the environment on NO and LRS, we ran additional models setting the variable representing the number of *fertile years* as the response variable. We found that it was also influenced by the early life environment and the environmental switch. Women born in the north-urban parish had more *fertile years* than those born in the rural parishes, which aligns with the result of earlier AFR for urban-born women. Similarly, an inverse relationship between a longer reproductive life span and an earlier age at maturity was also found in red deer females, leading to improvement of lifetime reproductive success [83]. We also observed that women who remained in the same urbanity had significantly more *fertile years* than women who switched from rural to urban, which would support the PAR hypothesis (Table 1). The large part of the variance captured by *fertile years* in the model for NO could explain why we do not observe a direct effect of the urbanity of birth or the switch in urbanity on NO. Therefore, the results for *fertile years* imply further support for the PAR hypothesis and provide additional information on the indirect effects of urbanity on NO.

## Conclusion

This study investigates the influence of environments at different life stages on preindustrial women reproductive performance. To do so, rather than analysing variations in environmental conditions across different time periods, we used information on the urbanity and the shore of the early life environment as well as the environmental switch between early life and adulthood to characterise environmental conditions experienced by women who had a complete reproductive history. We were thus able to separate the environmental effects attributed to the early life environment and those caused by the adult life environment, contributing towards a more comprehensive understanding of the relationship between environmental factors and reproductive outcomes. Our findings highlight the importance of both early and adult life environments on reproductive performance in humans, showing the relevance of the silver spoon hypothesis in preindustrial human populations but also finding evidence in favour of the PAR hypothesis and the importance of the quality of the adulthood environment in certain cases. The insights brought by our study on the impact of the environment on reproductive performance in humans has potential implications for various fields, including public health and evolutionary biology.

## Acknowledgments

We extend our sincere appreciation to the demographers, genealogists, and computer programmers, both past and present, who have dedicated their efforts to RPQA. In addition, we would like to thank Guillaume Blanchet and Dany Garant for their support and expertise, which have improved the manuscript.

## Supporting information captions

***Appendix S1: Additional analyses***

***Appendix S2: Sensitivity analysis***

***Figure S1: Geographical location of the 166 parishes of Nouvelle-France, established between 1616 and 1799.*** *Panel A shows the location of the parishes along the St Lawrence Valley, panel B shows a zoom into the area near Québec City and panel C shows a zoom into the area near Montréal. Conditions in the rural parishes were different from those in Québec City and Montréal, and conditions on the north shore (points in red) were also different from those in the south (points in blue). Conditions in the two cities were also different from each other*.

***Figure S2: Summary of the filters applied on the original dataset (N= 448,501) to obtain the subset used to perform the analyses shown in the manuscript (N= 7,203).*** *It also indicates which subset was used to calculate the number of offspring (NO) and lifetime reproductive success (LRS)*.

***Figure S3: Early life environmental effects on the Fertile Years (Panel A) and the Proportion between LRS and NO (panel B) according to the environment of birth, given by the urbanity (Rural or Urban) and the shore (North and South).*** *Rural and South are considered good environments and Urban and North are considered bad environments. The dots are the predicted marginal values, and the lines are their confidence intervals. Estimates with different letters are statistically different, given by a post-hoc Tukey’s pairwise multiple comparison test (P <0.05)*.

***Figure S4: Adult life environmental effects on the Fertile years (Panel A) and the Proportion between LRS and NO (Panel B and C) according to the switch in urbanity and to the switch in shore.*** *Staying in the same urbanity or in the same shore is considered as staying under the same conditions, while switching from urban to rural or north to south is seen as going from a bad to a good environment, and switching from rural to urban or south to north is considered going from a good to a bad environment. The dots are the predicted marginal values, and the lines are their confidence intervals. Estimates with different letters are statistically different, given by a post-hoc Tukey’s pairwise multiple comparison test (P <0.05)*.

***Figure S5: Early life environmental effects (Panel A) according to the environment of birth, given by the urbanity (Rural or Urban) and the shore (North and South) and adult life environmental effects (Panel B), according to the interaction between the switch in urbanity and the switch in shore, on Age at Marriage.*** *Rural and South are considered good environments and Urban and North are considered bad environments. Staying in the same urbanity or in the same shore is considered as staying under the same conditions, while switching from urban to rural or north to south is seen as going from a bad to a good environment, and switching from rural to urban or south to north is considered going from a good to a bad environment. The dots are the predicted marginal values, and the lines are their confidence intervals. Estimates with different letters are statistically different, given by a post-hoc Tukey’s pairwise multiple comparison test (P <0.05)*.

***Figure S6: Effects of the distance between the parish of birth and the parish of first reproduction on reproductive performance, given by Age at First Reproduction (Panel A), the Proportion between LRS and NO (Panel B) the Age at Marriage (Panel C).*** *The blue lines are the predicted marginal values, and the shaded areas describe the confidence interval*.

***Figure S7: Correlation between reproductive performance and reproductive schedule, for N=7,203.*** *Panel A shows the relationship between the age at first reproduction (AFR) in years and the number of offspring (NO) in terms of children born. Panel B displays the correlation between AFR and lifetime reproductive success (LRS) in terms of the number of children who survived to adulthood. Panel C illustrates the correlation between the number of offspring (NO) and the fertile years in years. Finally, Panel D exhibits the correlation between the fertile years and lifetime reproductive success in terms of the number of children who survived to adulthood*.

***Table S1: Descriptive statistics of the means of all the reproductive traits analysed.*** *They are calculated for N= 7,203, except Lifetime Reproductive Success which is calculated for N= 3,959*.

***Table S2: Descriptive statistics on the distribution of the population according to the environment of birth, for N= 7,203*.**

***Table S3: Descriptive statistics on the distribution of the population according to the environmental switch, for N= 7,203*.**

***Table S4: Output of the final model selected by stepwise backwards selection on a full model with all the variables present in the first column, for Fertile Years, the Proportion between Lifetime Reproductive Success (LRS) and Number of Offspring (NO) and the Age at Marriage*.**

***Table S5: Contributions of each fixed effect obtained via the ‘glmm.hp’ package and calculated by hierarchical partitioning of the marginal R² for generalised mixed-effect model, based on the output of r.squaredGLMM() from the ‘MuMIn’ package***.

***Code S1: Models***

***Code S2: Partitioned R***

Code S3: Plots

